# An Apoptotic Failsafe Program Involving PDGF-dependent PI 3- Kinase Activity Contributes to Plaque Composition in Atherosclerosis

**DOI:** 10.64898/2026.10.26.761862

**Authors:** Marius Vantler, Eva M. Berghausen, Moritz M. Schwab, Mario Zierden, Wolfram-Hubertus Zimmermann, Norbert Gerdes, Michelle Tallquist, Joachim Herz, Holger Winkels, Stephan Baldus, Stephan Rosenkranz

**Author notes:** **Address for correspondence:** Stephan Rosenkranz, MD Klinik III für Innere Medizin Herzzentrum der Universität zu Köln Kerpener Str. 62 D-50937 Köln, Germany.

## Abstract

**Background:** Growth and stability of atherosclerotic plaques are impacted by vascular smooth muscle cell (VSMC) apoptosis which is believed to be suppressed by growth factor dependent activation of phosphoinositide 3′-kinase (PI3K). Here, we analyzed the effects of chronically elevated platelet-derived growth factor receptor β (PDGFRβ) signaling on VSMC apoptosis and aortic plaque composition during atherogenesis.

**Methods:** We assessed atherosclerosis in low-density lipoprotein receptor deficient mice, lacking low-density lipoprotein receptor-related protein 1 in VSMCs (*LDLR^-/-^ smLRP1^-/-^* mice), in that smLRP1 deficiency resulted in PDGFRB overexpression. Apoptosis and downstream signaling was investigated in cultured aortic VSMCs.

**Results:** Chronically elevated PDGFRβ signaling in *LDLR^-/-^smLRP1^-/-^* mice unexpectedly increased VSMC apoptosis in atherosclerotic plaques. In vitro, PDGF exerted transient anti-apoptotic effects on VSMCs, whereas long-term stimulation promoted cell death. Mutation of the PI3K binding site in the PDGFRβ (PDGFRβ- F2/F2) reduced VSMC apoptosis in plaques as well as in cultured VSMCs. Further analyses revealed that PI3K isoform p110α mediates VSMC survival, whereas p110δ and p110β rather promote apoptosis. We identified a pro-apoptotic PDGFRβ-driven signaling pathway in VSMCs involving PI3K-dependent induction of c-Myc that in turn suppresses the expression of anti-apoptotic Bcl-2. PDGF-dependent downregulation of Bcl-2 increases the susceptibility of VSMCs to apoptotic agonists like H_2_O_2_ or Fas ligand. Consistently, Bcl-2 expression was elevated in plaques from PDGFRβ-F2/F2 mice.

**Conclusions:** Our study expands the understanding of the role of growth factors in regulating apoptosis and argues against their role as pure survival factors. This has direct implications in atherogenesis, as chronically elevated PDGF does not mediate VSMC survival in atherosclerotic plaques, but rather promotes cell death in the VSMC-rich fibrous cap, thus promoting plaque instability and rupture.

## INTRODUCTION

Numerous studies have demonstrated a protective role of VSMCs in atherosclerotic plaques.^1,2^ Advanced plaques comprise a fibrous cap mainly composed of VSMCs and extracellular matrix proteins overlying a lipid- and macrophage-rich necrotic core. Consequently, induction of VSMC apoptosis in atherosclerosis prone *ApoE*^-/-^ mice destabilizes the cap and predisposes the plaque to rupture^3,4^. Furthermore, chronic VSMC apoptosis leads to enhanced plaque growth, Marfan-like syndrome, and calcification.^3,4^ In humans, increased levels of VSMC apoptosis are seen in mature plaques compared with undiseased vessels,^5^ and plaques from patients with unstable symptoms show higher levels of apoptosis than those with stable lesions.^6,7^ Apoptosis is modulated by extracellular anti- and pro-apoptotic stimuli which regulate the expression and activity of apoptosis-associated proteins like caspases or Bcl-2 proteins.^8–10^ In atherogenesis, VSMC apoptosis is induced via death receptors like Fas^11,12^, oxidative stress^13,14^, pro-inflammatory cytokines, oxidized low-density lipoprotein (oxLDL), high levels of nitric oxide (NO), and mechanical injury.^6^ In contrast, receptor tyrosine kinases such as the platelet-derived growth factor receptor (PDGFRβ) or the insulin-like growth factor receptor are considered as potent inhibitors of cell death^15,16^ and are therefore believed to sustain cap thickness and plaque stability.^1,17^

The PDGFRβ is highly expressed in atherosclerotic lesions and assigned a key role in atherogenesis.^1,18^ Studies in atherosclerotic mice lacking the PDGFβ receptor in VSMCs indicate that PDGF is a key mitogen that promotes long-term plaque stability.^1,19^ Upon ligand binding, it recruits and activates cytoplasmic signaling molecules including Src family kinases (Src), phosphoinositide 3′-kinase (PI3K), and phospholipase Cγ (PLCγ) promoting cell migration, proliferation, and phenotypic switching.^20,21^ We previously demonstrated that *in vitro*, PDGF mediates anti-apoptotic signals in VSMCs exclusively via PI3K.^15^ Downstream of PDGFRβ, PI3K signaling is mediated by class IA PI3K isoforms, consisting of a regulatory p85 subunit and one of three distinct 110-kDa catalytic subunits (p110α, p110β, p110δ). Class IA PI3Ks are lipid kinases generating phosphatidyinositol-3,4,5-trisphosphate (PIP_3_),^22–24^ which serve as a docking site for downstream signaling molecules such as AKT, which is considered the major transducer of PI3K signal relay. AKT phosphorylates and thus inactivates proapoptotic proteins such as BAD, FoxO transcription factors, NFκB, caspase-9, and glycogen synthase kinase-3β.^15–17,25–27^

Despite these established signalling mechanisms, the pathobiology of plaque development and progression during atherogenesis is complex. Enhanced PDGFRβ/PI3K signaling in VSMCs via smLRP1 knockout in LDLR^-/-^ mice resulted in phenotypes which are attributed to increased VSMC apoptosis, including Marfan-like syndrome and accelerated plaque growth.^3,4,28,29^ Hence, these data challenge the concept of PDGF as a pure survival factor, particularly, when PDGFRβ/PI3K signaling is chronically elevated as in atherogenesis. Here, we explored the impact of chronic PDGF signalling via PI3K isoforms and its impact on aortic atherosclerotic plaque composition in detail.

## METHODS

### Generation of mouse strains & aorta preparation

Generation of smLRP1^-/-^; LDLR^-/-^ and smLRP1^-/-^; LDLR^-/-^; PDGFRβ-F2/F2 mice and aorta preparation were described elsewhere^18,29^. Briefly, smooth muscle-specific knock out of LRP1 and LDLR were achieved using the Cre / *loxP* system. All animals were maintained on standard rodent chow diet. Hearts from 12 month old mice were removed including the root of the aortic arch and preserved in 4 % paraformaldehyde (PFA) for histological studies.

### TUNEL assay and immunohistochemistry

Paraffinsections from the aortic arch were deparaffinated and stained with hematoxylin and eosin. Apoptotic cells were detected using a DeadEnd™ Fluorimetric TUNEL System assay (Promega) according to the manufacturerś instructions. Afterwards, sections were blocked (10% (w/v) nonfat dry milk in PBS), and stained with a mouse monoclonal, Cy3^™^-conjugated α-smooth muscle actin antibody (Sigma Aldrich; 1:200) for 1 h. The fluorescein-12-dUTP-labeled DNA and Cy3^™^-stained cells were visualized by fluorescence microscopy using a NIKON Eclipse E800 microscope equipped with a NIKON DMZ1200 camera. Apoptotic cells were analyzed after performing overlay projections. The percentage of TUNEL positive stained cells was determined by counting the number of TUNEL positive cells and DAPI stained cells in Cy3^™^-stained areas of the plaque (5 cross-sections from each mouse strain (n = 6, respectively)). Bcl-2 was stained using a polyclonal Bcl-2 antibody (Santa Cruz, 1:100). Cap and plaque areas were determined using imageJ analysis software (National Institutes of Health, Bethesda, Maryland).

### Cell Culture

VSMC were isolated from rat thoracic aorta (Wistar Kyoto; 6-10 weeks old; Charles River Wega GmbH, Sulzfeld, Germany) by enzymatic dispersion as described ^30^. Cells were grown in a 5 % CO_2_ atmosphere at 37 °C in Dulbecco’s modified Eagle’s medium (DMEM) supplemented with 100 U/ml penicillin, 100 µg/ml streptomycin, and 10 % FCS. ChiR mutants and their maintenance were previously described.^15,31^

### Induction and detection of apoptosis

A Cell Death Detection ELISA (Roche Diagnostics) was used to quantify apoptosis by the detection of cytoplasmic histone-associated DNA fragments (mono- and oligonucleosomes) according to the manufacturerś instructions. VSMCs (5,000 cells/well) were plated into 96-well plates in DMEM containing 10% FCS, and starved for 24 h in DMEM. To assess the anti-apoptotic effect of PDGF (50 ng/ml), the cells were treated with H_2_O_2_ (100 µM) or UV-irradiated (254 nm, 70 mJ/cm^2^) and subsequently cultured for 2 to 16 h in the absence or presence of ligand. To analyze the pro-apoptotic effect of PDGF, starved VSMCs were pre-stimulated for 24 h with PDGF and subsequently treated for additional 16 h with H_2_O_2_ (50 or 100 µM). Where indicated, VSMC were transfected with Bcl-2 siRNA or non-silencing (ns) siRNA 24 h prior to starvation. Alternatively, starved VSMC were stimulated for 24 h to 96 h with PDGF or M-CSF. The cell lysates were collected, and apoptosis was measured using the Cell Death Detection ELISA.

### xCelligence assay

The xCelligence assay is based on electrical impedance readings in cell monolayers plated in wells containing built-in gold electrodes.^32^ We have used the analyzer, 96 well e-plates, and the integrated software from Roche Diagnostics GmbH. Cells were plated at a density of 5,000 cells/well in 200 μl of DMEM containing 10 % FCS, and then the plate was installed on the analyzer. The analyzer and the installed plate were placed in a standard cell culture incubator, at 37 °C in a humidified atmosphere of 5 % CO_2_. Cells were allowed to adhere to plates overnight, starved for 24 h in DMEM, and then stimulated for 96 h with FCS (10 %) or FCS and PDGF (50 ng/ml). The analyzer was programmed to take readings during the whole cell culturing period. Data were recorded and analyzed using the integrated software.

### Flow cytometric apoptosis detection

To distinguish between healthy and apopototic cells by detecting changes in the mitochondrial transmembrane potential, VSMC were stained with the MitoCapture dye (Mitochondrial Apoptosis Staining Kit, PromoKine) according to manufacturerś instructions after exclusion of nonviable cells via a forward and side scatter gate. Flow cytometry was performed on a FACSCalibur and analyzed by CellQuest Pro software (BD Biosciences).

### Transfection with siRNA

VSMC grown to 30% confluency in 96-well plates were transfected with 10 nM siRNAs specific for Bcl-2, ERK, or a non-silencing control (Qiagen) with Lipofectamine RNAiMax (Invitrogen) as described by the manufacturer. Apoptosis assays were performed 72 h later as described above.

### Western blot Analysis

Quiescent VSMC were left resting or stimulated with 50 ng/ml PDGF-BB or M-CSF for times indicated in the presence or absence of inhibitors as indicated. Cells were washed twice with HS (20 mM Hepes, pH 7.4, 150 mM NaCl) and then lysed in EB (10 mM Tris-HCl, pH 7.4, 5 mM EDTA, 50 mM NaCl, 50 mM NaF, 1% Triton X-100, 0.1 %

BSA, 20 µg/ml aprotinin, 2 mM Na_3_VO_4_, 1 mM phenylmethylsulfonyl fluoride). Lysates representing approximately 3x10^6^ cells were centrifuged (20 min, 12,000 × g), and the supernatants were resolved on a SDS-PAGE gel and probed with antibodies raised against Bcl-2, Bak, Bax, Bid, Bim, Noxa, phosphorylated Akt, and RasGAP (lysate control).

### RNA isolation, reverse transcription and quantitative real-time PCR

RNA was isolated using Trizol (Invitrogen) and quantified by spectrophotometry. 2 µg of total RNA was transcribed with SuperScriptII RT (Invitrogen). Individual samples of 20 ng cDNA were subjected to PCR amplification with AmpliTaqGold Polymerase II (Applied Biosystems) utilizing gene-specific primers and fluorogenic probes (5’ FAM and 3’ TAMRA; Supplementary Table) in an ABI PRISM 7900HT Sequence Detection System (Applied Biosystems). The PCR conditions are as follows: 30 sec at 50 °C and 10 min at 95 °C followed by 45 cycles of 30 sec at 95 °C and 1 min at 58 °C. A template-free negative control was included in each experiment. GAPDH expression was similar in all study groups and was therefore employed to normalize for differences in RNA quantity and RT-efficiency. Quantification of transcripts was performed using the 2^-ΔΔCt^ method.^33^

### Materials

PDGF-BB and M-CSF were purchased from Promo Cell (Heidelberg, Germany). The antibody against RasGAP (69.3) was a kind gift from Andrius Kazlauskas and Alex Toker (Harvard Medical School, Boston, USA). The polyclonal Phospho-AKT (Ser473), was from Cell signaling technologies (Beverly, USA). The monoclonal Bcl-2, Bak, Bax,

Bid, Bim, and Noxa antibodies were from Santa Cruz Biotechnology, Inc..^34^ Specific pharmacological inhibitors were from Calbiochem (Darmstadt, Germany).

## Statistical analysis

Data are presented as mean±SD. Two-group comparisons used 2-tailed Student t test; 1-way ANOVA or 2-way ANOVA were used for multiple group comparisons; P<0.05 was considered significant. P values follow AHA formatting guidelines.

## RESULTS

### Chronic PDGFRβ/PI3K signaling exerts pro-apoptotic effects in atherosclerotic plaques

We analysed the effects of chronic PDGFRβ/PI3K signaling on VSMC apoptosis in atherosclerotic plaques by means of LDLR^-/-^smLRP1^-/-^ and PI3K-binding deficient LDLR^-/-^smLRP1^-/-^PDGFRβ-F2/F2 mice. Lack of LRP1 expression resulted in elevated PDGFRβ expression and activation in both LDLR^-/-^smLRP1^-/-^ and LDLR^-/-^smLRP1^-/-^ PDGFRβ-F2/F2 mice compared to LDLR^-/-^ control mice by mechanisms that are not fully understood (**Figure 1A)**^18^. Quantification of apoptotic TUNEL positive cells in the VSMC-rich fibrous cap (**Figure 1B**) revealed increased apoptosis in LDLR^-/-^smLRP1^-/-^ mice (2.9±2.3 % vs. 0.7±0.9 % in LDLR^-/-^ mice) (**Figure 1C**). Deletion of the PI3K binding site in the PDGFRβ in LDLR^-/-^smLRP1^-/-^PDGFRβ-F2/F2 mice significantly reduced this effect (2.0±1.9 % TUNEL positive cells), suggesting that increased expression and activity of the PDGFRβ exerts pro-apoptotic properties in VSMCs via activation of PI3K.

**Figure 1.**
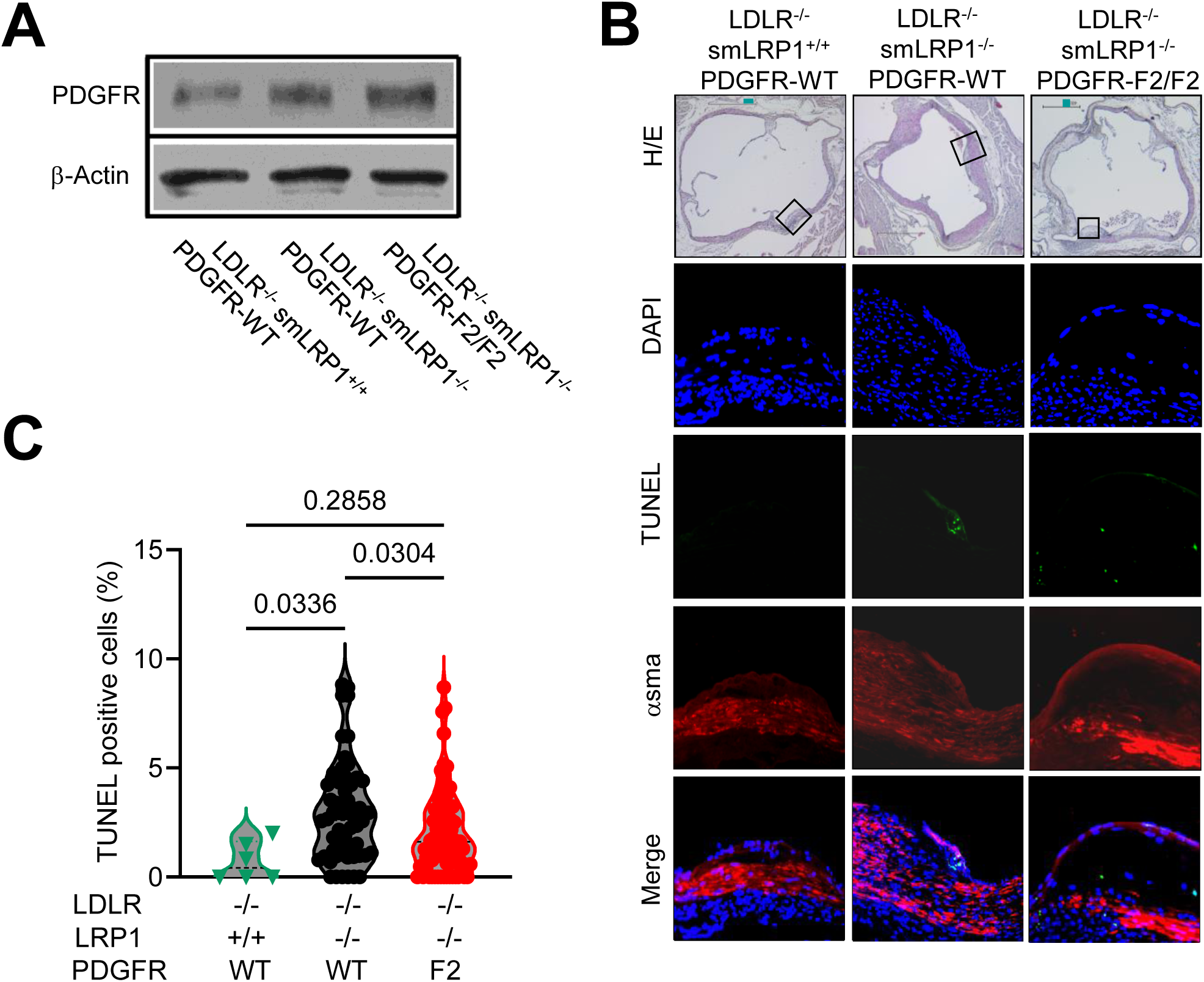
PDGFRβ- and PI3K-dependent increase of VSMC apoptosis in atherosclerotic plaques. (**A**) Protein amounts of the PDGFRβ in VSMCs from LDLR^-/-^ smLRP1^+/+^PDGFRβ control mice, LDLR^-/-^smLRP1^-/-^PDGFRβ mice, and LDLR^-/-^ smLRP1^-/-^PDGFRβ-F2/F2 mice. The β-actin immunoblot is shown as a loading control. (B) Hematoxylin and eosin (H&E), DAPI, TUNEL, α-sm-actin stainings and merge of lesions from indicated mice. (**C**) Quantitative analysis of TUNEL stainings from indicated sections (LDLR^-/-^smLRP^+/+^PDGFRβ (n=6 sections from 1 animal), LDLR^-/-^ smLRP1^-/-^PDGFRβ (58 sections from 6 animals) and LDLR^-/-^smLRP1^-/-^PDGFRβ-F2/F2 mice (81 sections from 6 animals)). Data is presented as means ± SD. *P* values were determined by one-way ANOVA followed by Tuckeýs multiple comparison.

### Differential regulation of VSMC apoptosis by PDGF/PI3K signaling

This unexpected finding is in contrast to previous *in vitro* data from our lab demonstrating that PDGF protects VSMCs against UV-irradiation induced apoptosis via activation of PI3K/AKT^15^. However, analysis of VSMC apoptosis and AKT activation at different time points following PDGF stimulation revealed that the anti-apoptotic effect of PDGF/PI3K/AKT appears to be only transient. No significant protection against apoptosis and activation of AKT were detected 16 h after PDGF stimulation (**Figure 2** **A,B**). Therefore, we next explored PDGF long-term effects on VSMC apoptosis beyond 16 h. We pre-stimulated VSMCs for 24 h with PDGF and subsequently induced apoptosis with H_2_O_2_. PDGF pre-stimulation caused a marked increase of cell death (5.7±1.7 fold increase over control) (**Figure 2C**). This long-term effect was maximal using 100 ng/ml PDGF (**Figure S1**). Additionally, pre-stimulation with PDGF significantly enhanced Fas ligand (FasL) mediated apoptosis (**Figure 2D**). Furthermore, we analyzed mitochondrial dysfunction in VSMCs, a hallmark of apoptotic pathways which are mainly controlled by Bcl-2 proteins. Flow cytometric analysis revealed that 23.5±7.5 % of untreated control cells contained dysfunctional mitochondria. Induction of apoptosis by H_2_O_2_ alone increased this cell fraction to 33.3±1.6 %, whereas pre-stimulation with PDGF further enhanced the percentage of VSMCs comprising dysfunctional mitochondria to 59.8±8.0 % (**Figure 2E,F**).

**Figure 2.**
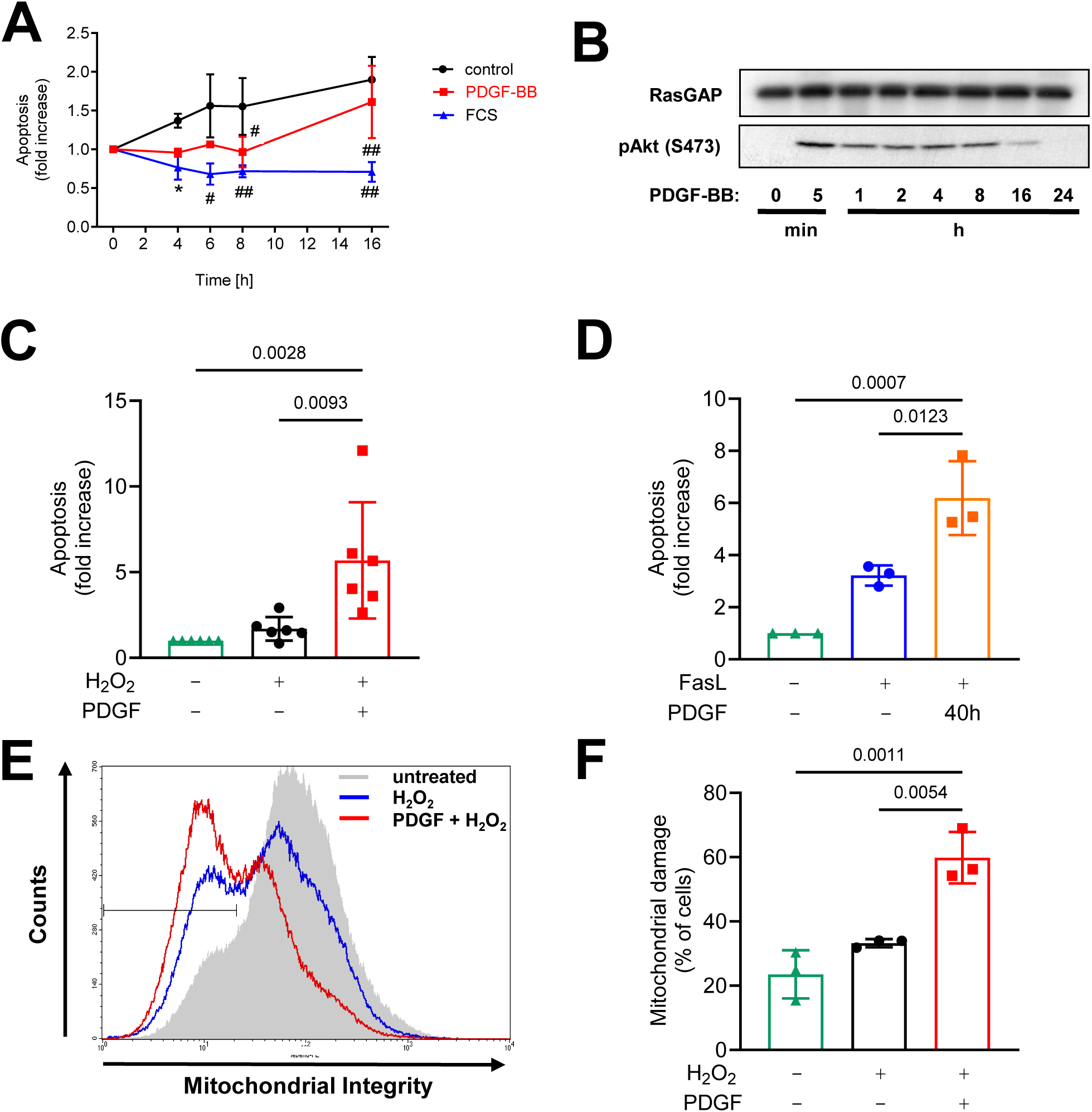
PDGF-mediated VSMC survival is only transient, and long-term PDGF-BB stimulation increases the susceptibility to apoptosis. (**A**) DNA fragmentation was assessed at the indicated time points after UV irradiation. VSMC were stimulated with PDGF-BB (conc) or FCS (conc) immediately following UV irradiation (n=3). Data are presented as means ± SD. *P* values were determined by one-way ANOVA followed by Tuckeýs multiple comparison (\**p*<0.05 versus control; ^#^*p*<0.001 versus control; ^##^*p*<0.0001 versus control or versus PDGF (16h)). (**B**) Starved VSMC were stimulated for the indicated periods of time with PDGF (50 ng/ml) and lysates were probed with a phospho-specific AKT antibody. RasGAP blot is shown as a loading control. (**C**) Apoptosis of cell stimulated with H_2_O_2_ (100 µM) (n=6) or (**D**) FasL (100 ng/ml) (n=3) in the absence or presence of PDGF-BB (50 ng/ml, 24h pre-stimulation) as quantified by nucleosomal ELISA. (**E** and **F**) Flow cytometric analysis of mitochondrial damage of untreated, H_2_O_2_ (50 µM) stimulated, or PDGF and H_2_O_2_ stimulated cells (n=3). Data is presented as means ± SD. *P* values were determined by one-way ANOVA followed by Tuckeýs multiple comparison.

We next explored whether PDGF is able to induce apoptosis by itself. Therefore, we stimulated VSMCs for up to 4 days with either FCS (10%), PDGF, or left them untreated. While FCS protected against apoptosis throughout the observation period, chronic stimulation with PDGF for 4 days led to a modest but significant increase of apoptosis (1.9±0.2-fold) (**Figure 3A**). Interestingly, this effect was even more pronounced when PDGF and FCS were simultaneously applied to the cells (8.9±4.7- fold at day 4).

**Figure 3.**
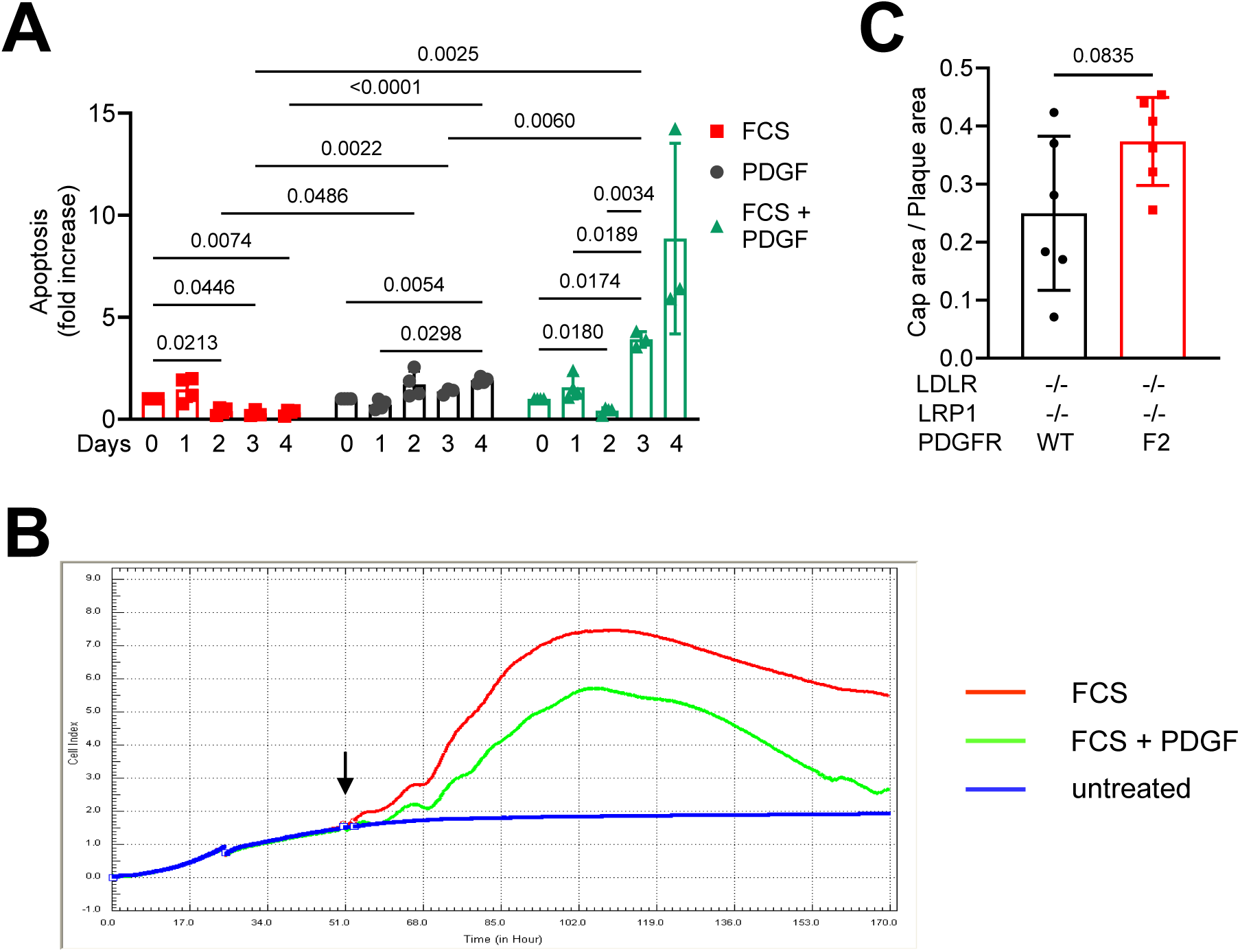
Chronic stimulation with PDGF induces VSMC apoptosis (**A**) Apoptosis of VSMCs treated with FCS (10%), PDGF (50 ng/ml), a combination of both, or left untreated for 1, 2, 3, or 4 days (n=3-4). Error bars are means ± SD. *P* values were determined by two-way ANOVA followed by Tuckeýs multiple comparison. (**B**) Dynamic monitoring of VSMC confluency over 170 h in non-stimulated (blue line), FCS (red line), or PDGF + FCS (green line) stimulated cells. Stimulation with FCS or FCS + PDGF is indicated by an arrow. (**C**) Cap area to plaque area ratio of plaques from LDLR^-/-^smLRP1^-/-^PDGFRβ (n=15 plaques from 6 animals) and LDLR^-/-^smLRP1^-/-^ PDGFRβ F2/F2 mice (6 animals, respectively); *P* value was determined by Welch′s *t* test.

### Impact of PDGF and FCS on cell-substance impedance over time

In order to investigate whether this robust increase of apoptosis overruns PDGF and FCS induced cellular proliferation ultimately leading to a decrease in cell number, we measured changes of cell confluence in real-time. To this end, we detected alterations in the cell-substrate impedance (displayed as “cell index”) depending on the cell number in the culture dish.^32^ After an initial increase of the cell index due to PDGF and FCS mediated VSMC proliferation, a decrease of the cell index was observed two days after stimulation with FCS and PDGF (**Figure 3B**) as compared to cells stimulated with FCS alone. Four days after PDGF/FCS stimulation, the cell index nearly reached the same level than unstimulated cells, suggesting that PDGF- and FCS-induced apoptosis led to a decrease in cell number. PDGF-induced proliferation and migration which is mainly mediated by PI3K^21,29^ contribute to development and stabilization of the fibrous cap. **Figure 3C** demonstrates that abrogation of PDGFRβ-induced activation of PI3K did not thin out the cap region in plaques from LDLR^-/-^smLRP1^-/-^ PDGFRβ-F2/F2 mice, but rather increased the cap area (0.37±0.08 vs. 0.25±0.13 in LDLR^-/-^smLRP1^-/-^ mice). These data indicate that PDGFRβ-mediated pro-apoptotic signaling can compensate for or even exceed migratory and proliferative responses *in vitro* and *in vivo*.

### Systematic evaluation of the role of PDGFRβ-associated signaling molecules

To characterize the impact of each PDGFRβ-associated signaling molecules (Src, PI3K, RasGAP, SHP-2, and PLCγ) on the pro-apoptotic effect induced by long-term PDGFRβ activation, we took advantage of chimeric c-fms/PDGFRβ (ChiR) mutants^15^ (**Figure S2**). These mutants are not compromised by short half-lives like many inhibitors and are therefore most suitable to investigate long-term PDGFRβ mediated effects. ChiR mutants contain the ligand binding domain of c-fms and are activated by M-CSF. In the cytoplasmic domain of the PDGFRβ, specific binding sites were individually deleted by tyrosine to phenylalanine mutations. We stimulated each ChiR mutant for 4 days with M-CSF. Activation of ChiR-WT induced apoptosis to comparable extent as the endogenous PDGFRβ (2.6±0.3-fold) (**Figure 4A**). Analysis of the *subtraction panel* which contains mutants that lack one specific binding site, respectively, demonstrated that abrogation of PI3K binding (F40/51) completely abolished M-CSF-induced apoptosis whereas the lack of either Src, SHP-2 or PLCγ binding had no influence (**Figure 4A**). Interestingly, activation of the F771 mutant which lacks the binding site for the negative regulator of Ras, RasGAP, mediated a much more pronounced increase of apoptosis than the ChiR-WT (7.8±1.3-fold). This was expected since activation of Ras is necessary to fully activate PI3K. The characterization of the *add-back* mutants which bind only one signaling molecule in addition to Src (which is required for ligand-induced receptor activation), respectively, revealed that activation of PI3K via the Y40/51 receptor was sufficient to mediate the same pro-apoptotic effect than the ChiR-WT (**Figure 4B**). These findings are consistent with the *in vivo* data which suggest that deletion of the PI3K binding site in the PDGFRβ polypeptide reduces apoptosis in atherosclerotic plaques.

**Figure 4.**
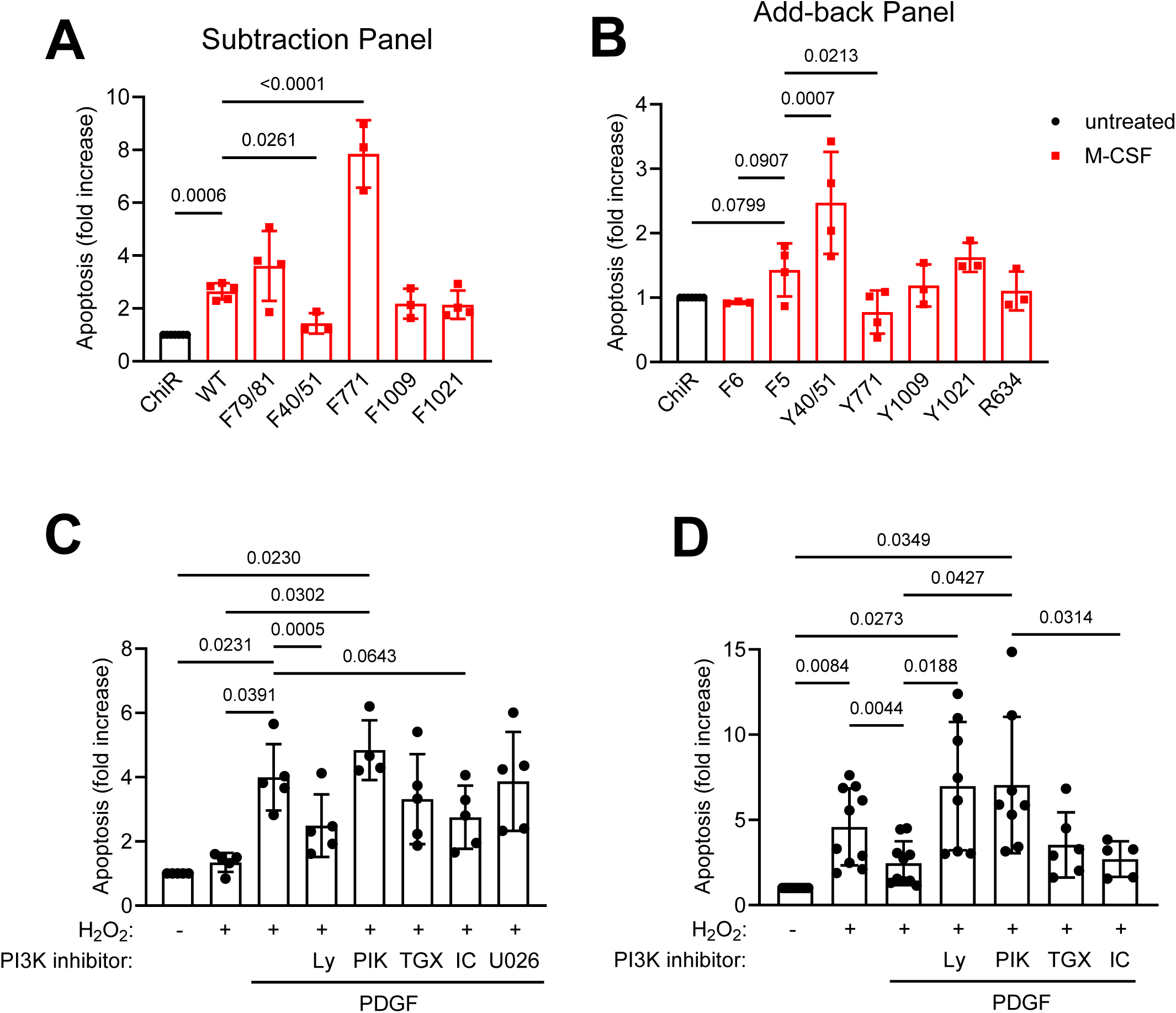
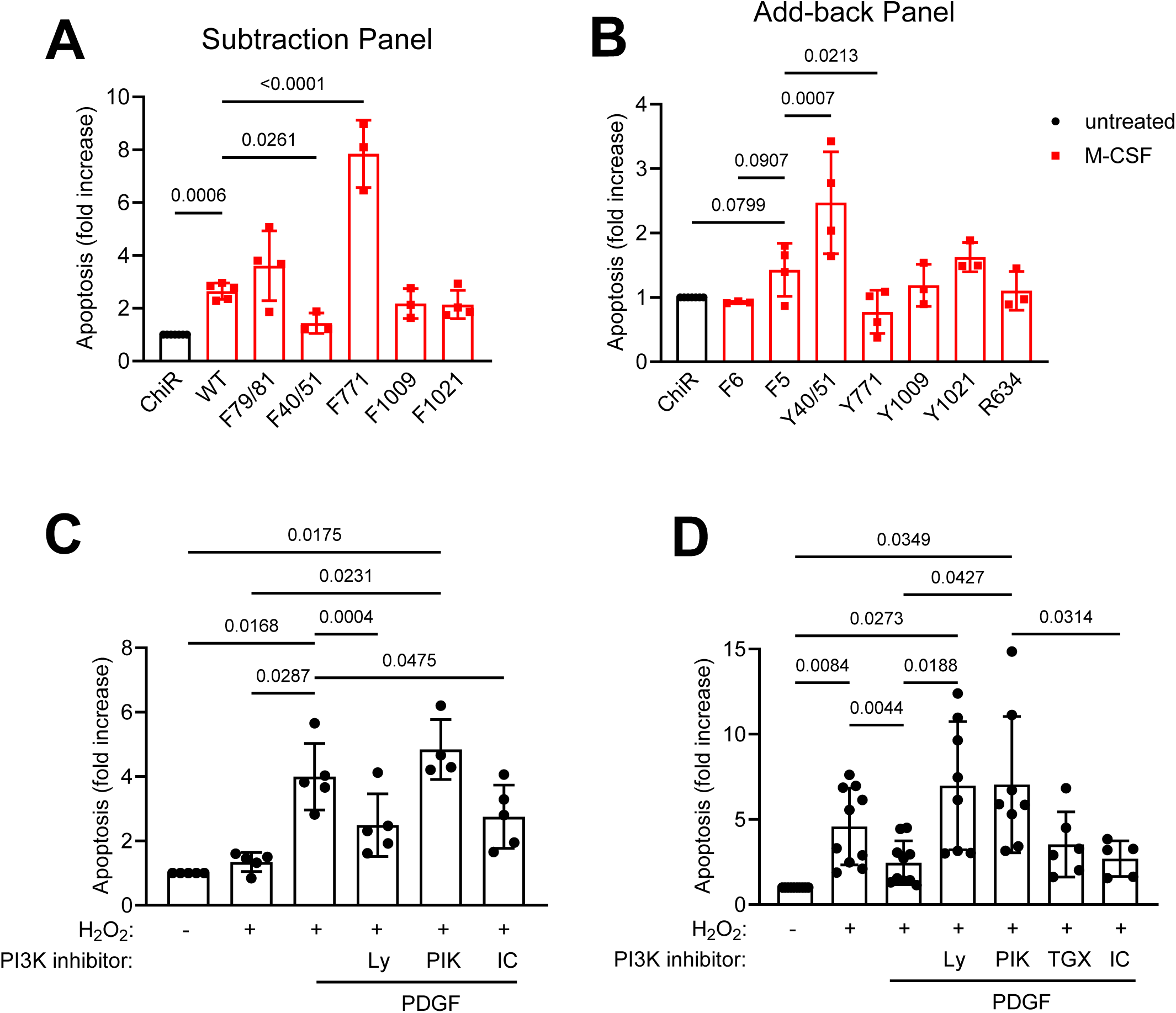
PDGFRβ-mediated apoptosis depends on PI3K. (**A**) Quantitative analysis of M-CSF-induced DNA fragmentation of VSMC expressing chimeric c-fms/PDGFRβ (ChiR) mutants from the subtraction panel (Supplementary figure 3a). Cells were stimulated for 4 days with M-CSF. (**B**) Quantitative analysis of M-CSF-induced DNA fragmentation of VSMC expressing ChiR from the add-back panel (see **Supplemental** Figure 3B). Data were measured in triplicates and represent means ± SD from at least 3 independent experiments. Data were normalized by the apoptosis rate of unstimulated cells in the absence of M-CSF for each mutant separately. The relative apoptosis rate for unstimulated cells in the absence of M-CSF is 1 for each ChiR and indicated as “ChiR”. Data is presented as means ± SD. *P* values were determined by one-way ANOVA followed by Tuckeýs multiple comparison. (**C**) Analysis of H_2_O_2_ induced DNA fragmentation in cells pretreated with PDGF for 24h prior to H_2_O_2_ addition in the presence or absence of inhibitors against PI3K (Ly-294002, 10 µM), p110α (PIK075, 300 nM), p110β (TGX-221, 1µM), p110δ (IC-87114, 1µM), or MEK1/2 (U0126, 10µM). n=3-6. (**D**) Analysis of apoptosis in cells simultaneously treated with PDGF and H_2_O_2_ (16h) in the presence or absence of indicated inhibitors (n=5-10). *P* values were determined by one-way repeated measure ANOVA followed by Tuckeýs multiple comparison.

### Differential regulation of apoptosis by catalytic class IA PI3K isoforms

Consistent with the above findings, utilization of the pan PI3K inhibitor LY-294002 *in vitro* demonstrated that the catalytic activity of PI3K is necessary for the PDGF-mediated enhancement of apoptosis. In contrast, inhibition of MEK1/2 did not exert any effects excluding a prominent role of the Ras/ERK pathway in this context (**Figure 4C**).

Furthermore, selective inhibition of catalytic PI3K subunits indicated isoform specific effects on PDGF mediated apoptosis. While selective inhibition of PI3K/p110α did not affect long-term PDGF dependent apoptotic effects on H_2_O_2_ treated cells, inhibition of p110δ and p110β rather ameliorated cell death (**Figure 4C**). In contrast, short-term survival effects by PDGF in VSMCs, which were simultaneously stimulated with PDGF and H_2_O_2_ for 16h, were abrogated by p110α inhibition, whereas p110δ and p110β inhibitors were ineffective (**Figure 4D**).

### Chronic PDGF stimulation induces pro-apoptotic effects via regulation of Bcl-2

The family of Bcl-2 proteins constitute central regulators of apoptosis. Pro-apoptotic Bcl-2 proteins increase the mitochondrial permeability by forming pores in the outer mitochondrial membrane, whereas anti-apoptotic Bcl-2 proteins bind and antagonize the pro-apoptotic ones. We analyzed protein expression of different Bcl-2 family members 8, 16, and 24 h following PDGF stimulation. Western blot analysis revealed that PDGF at 16 and 24 h downregulates anti-apoptotic Bcl-2, whereas the expression levels of pro-apoptotic Bak, Bax, Bid, Bim and Noxa remained unaltered (**Figure 5A**). Accordingly, the Bcl-2/Bax protein ratio decreased by ∼60 % (**Figure 5B**). Real-time PCR analysis revealed that the Bcl-2/Bax ratio declined at the mRNA level by ∼80 % (**Figure 5C**), indicating that PDGF regulates expression of Bcl-2 proteins at the transcription level. Furthermore, we did not detect the cleavage product of Bid (tBid) suggesting that death receptor pathways are not involved in the PDGF-induced pro-apoptotic response. Consistent with the pro-apoptotic long-term effect shown in **Figure 3A**, PDGF dependent downregulation of Bcl-2 was constant over the entire observation period of 4 days (**Figure 5D**). This effect appears to be PDGF-specific as chronic stimulation with FCS alone had only little influence on Bcl-2 expression, whereas the combination of FCS and PDGF efficiently downregulated Bcl-2 (**Figure 5D, Figure S3**). siRNA induced knockdown of Bcl-2 resulted in an enhancement of H_2_O_2_-induced apoptosis which was comparable to the PDGF-effect on H_2_O_2_-induced apoptosis in control siRNA transfected cells (**Figure 5E**). PDGF pretreatment only poorly enhanced H_2_O_2_-induced apoptosis in Bcl-2 knockdown cells, suggesting that downregulation of Bcl-2 is largely responsible for PDGF-mediated VSMC apoptosis. To verify this signaling pathway in atherosclerotic plaques, we analyzed Bcl-2 expression in LDLR^-/-^smLRP1^-/-^ mice as well as in LDLR^-/-^smLRP1^-/-^PDGFRβ F2/F2 mice, in which the PDGFRβ does not bind and activate PI3K. **Figure 5F** demonstrates that in smLRP1^-/-^mice harbouring the wild type PDGFRβ, Bcl-2 expression was suppressed both in the VSMC-rich fibrous cap and media when compared to PDGFRβ- F2/F2 expressing mice. These data indicate that upregulation of the PDGFRβ in plaque VSMCs leads to a PI3K-dependent downregulation of Bcl-2 and increased VSMC apoptosis in mice rather than to cellular survival.

**Figure 5.**
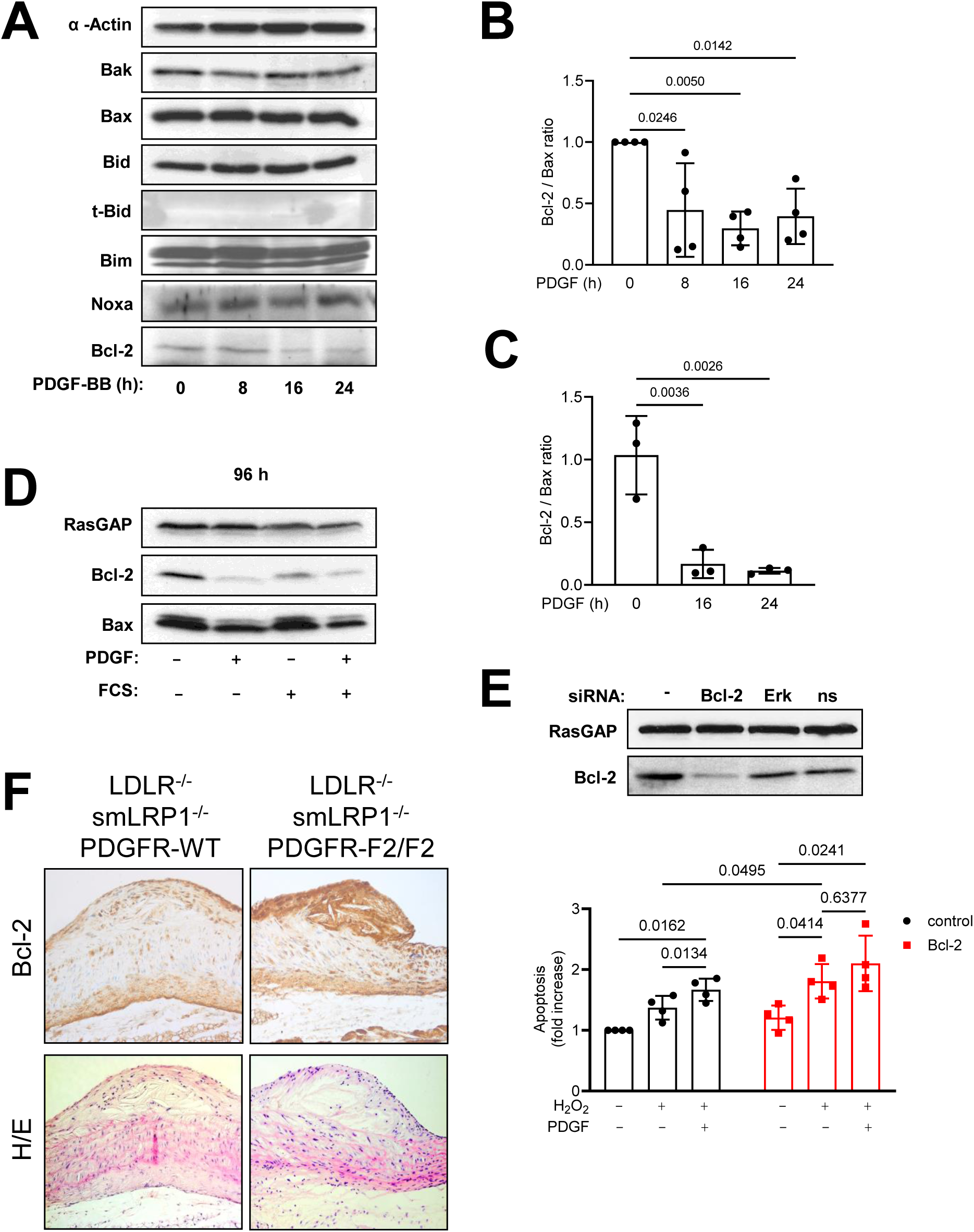
Chronic PDGF stimulation impairs Bcl-2 expression (**A**) Western blot analysis of Bcl-2 proteins expression in VSMCs stimulated for different periods of time with PDGF (left). Membranes were probed with antibodies against RasGAP (loading control) or indicated Bcl-2. proteins (**B**) Densitometrical analysis of the Bcl-2/Bax protein ratio (n=4). (**C**) Western blot analysis of cells treated for 4 days with PDGF and/or FCS or left untreated as indicated. (**D**) Ratio of Bcl-2 and Bax mRNA expression (n=3). *P* values were determined by one-way ANOVA followed by Tuckeys multiple comparison. (**E**) *Left*: Western blot analysis with a specific Bcl-2 antibody demonstrating siRNA mediated knockdown of Bcl-2. ERK and non-silencing (ns) siRNAs were used as negative controls. RasGAP blot is shown as a loading control. *Right*: Quantitative analysis of DNA fragmentation in control siRNA transfected or Bcl- 2 knockdown cells. Apoptosis was induced by H_2_O_2_ (50 µM) as indicated 24 h upon PDGF (50 ng/ml) stimulation (n=4). *P* values were determined by two-way ANOVA followed by Tuckeys multiple comparison. Data represent means ± SD (**F**) Immunostaining for Bcl-2 (top) and Hematoxylin and eosin (H&E) stainings in a representative atherosclerotic plaque from either LDLR^-/-^smLRP1^-/-^PDGFRβ mice (left), or LDLR^-/-^smLRP1^-/-^PDGFRβ-F2/F2 mice (right).

### PDGF/PI3K-dependent regulation of apoptosis involves c-Myc

Numerous studies demonstrated that c-Myc regulates the expression of Bcl-2 proteins.^35^ Furthermore, c-Myc is upregulated in human plaque VSMCs^36^. Here, we show that c-Myc is transiently induced by PDGF prior to Bcl-2 downregulation (**Figure 6A**). Knockdown of c-Myc prevented both PDGF induced downregulation of Bcl-2 (**Figure 6B**) and PDGF-induced increase of H_2_O_2_-mediated apoptosis (**Figure 6C**), indicating that it is involved in pro-apoptotic PDGFRβ signaling. To investigate the influence of PI3K on the expression of c-Myc and Bcl-2, we stimulated the ChiR-WT, F40/51, F5 and Y40/51 mutants for 3 and 24 h with M-CSF. Activation of PI3K alone (Y40/51) was sufficient to induce c-Myc and to downregulate Bcl-2 to the same extent as ChiR-WT, whereas mutation of the PI3K binding site (F40/51) completely abrogated these effects, as also seen in ChiR-F5 expressing cells (**Figure 6D**). These data demonstrate that PDGF-induced Bcl-2 downregulation and apoptosis is mediated through PI3K mediated induction of c-Myc.

**Figure 6.**
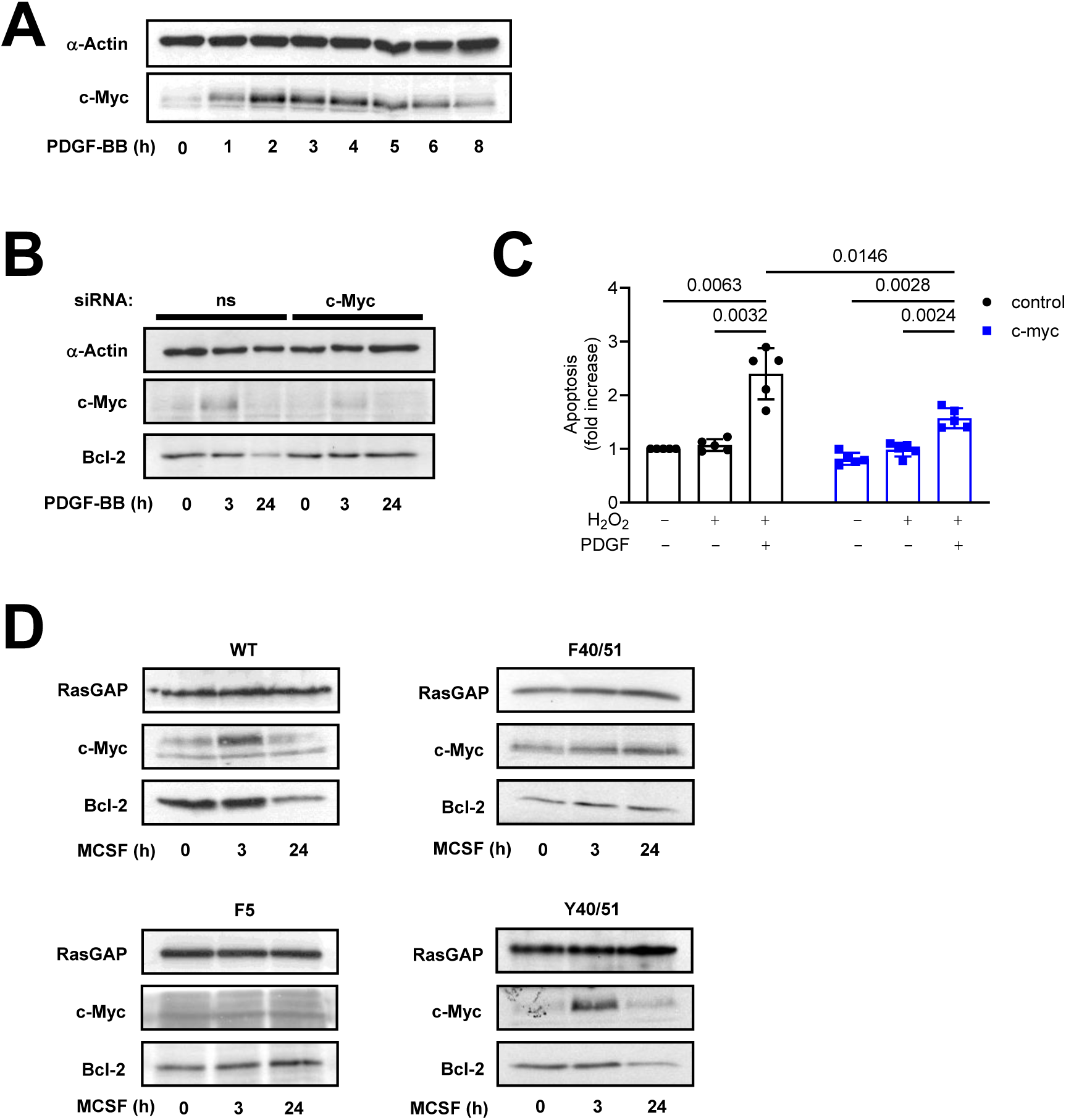
Stimulation with PDGF decreases Bcl-2 expression through PI3K dependent up-regulation of c-Myc. (**A**) PDGF transiently induces c-Myc in VSMC. Representative immunoblot with lysates from VSMCs treated for the indicated periods of time with PDGF. Membranes were probed with antibodies against RasGAP (loading control) or c-Myc. (**B**) Western blot analysis of c-Myc and Bcl-2 expression in control siRNA transfected cells or c-Myc knockdown cells. α-actin blot is shown as a loading control. (C) Apoptosis in control cells or c-Myc knockdown cells. VSMC were stimulated as indicated with H_2_O_2_ and/or PDGF. Data were measured in triplicates, normalized by the apoptosis rate of unstimulated cells which were transfected with non-silencing siRNA. Data from 5 independent experiments are shown as means ± SD. *P* values were determined by two-way ANOVA followed by Tuckeýs multiple comparison. (**D**) Western blot analysis of PDGFRβ mediated expression of c-Myc and Bcl-2 in cells expressing ChiR-WT, F40/51 which lacks only the PI3K binding site, F5 that lack all binding sites or Y40/51 which binds only PI3K. Cells were stimulated for the indicated periods of time with 50 ng/ml M-CSF.

## DISCUSSION

The data presented herein challenge the concept of PDGF being a pure survival factor and rather indicate that chronically elevated PDGF signaling may also exert pro-apoptotic effects in VSMCs that are relevant for atherosclerotic plaque composition and stability. The main results include: (i) Chronically enhanced PDGFRβ signaling in LDLR^-/-^smLRP1^-/-^ mice results in VSMC apoptosis in atherosclerotic plaques *in vivo*; (ii) This effect was blunted in animals expressing a PDGFRβ unable to bind PI3K (LDLR^-/-^smLRP1^-/-^PDGFRβ-F2/F2 mice); (iii) Chronic pro-apoptotic PDGFRβ signaling in VSMCs involves PI3K-dependent induction of c-Myc and subsequent suppression of anti-apoptotic Bcl-2; (iv) Pro- versus anti-apoptotic PDGFRβ signalling is differentially regulated via catalytic class IA PI3K isoforms.

Dysregulation of cellular apoptosis is a hallmark of many diseases like myocardial infarction or cancer. In atherogenesis, it was shown that apoptosis of plaque VSMCs causes a thinning of the fibrous cap and thereby predisposes plaques to rupture. Furthermore, VSMC apoptosis resulted in an accelerated plaque growth and Marfan-like syndrome.^3,4,6^ Most of these features are recapitulated in LDLR^-/-^smLRP1^-/-^ mice. smLRP1^-/-^ mice are characterized by elevated PDGF and transforming growth factor β (TGFβ) signaling^28^. Importantly, mutation of the PI3K binding site in the PDGFRβ polypeptide largely abolished plaque apoptosis, and as previously shown, accelerated plaque growth and Marfan-like syndrome.^29^ Accordingly, chronic mTOR activation downstream of PI3K/AKT induces a degradative VSMC phenotype.^37^ These data indicate that during elevated, chronic PDGFRβ activation, downstream PI3K activity is primarily responsible for these effects. Chronic PDGFRβ/PI3K activation does not only induce VSMC apoptosis as shown here, but also mediates VSMC proliferation and migration.^18,21,29,38^ In fact, depletion of PDGFRβ signaling in VSMCs resulted in decreased indices of lesion stability, indicating that PDGFRβ mediated VSMC proliferation and migration is required to maintain fibrous cap stability.^1,19^

While all of these in part opposing cellular responses contribute to plaque size and composition, their relative contribution appears crucial for plaque growth and composition. In cell culture, we were able to demonstrate that PDGF driven apoptosis can overrun PDGF mediated VSMC proliferation, and in LDLR-/-smLRP1-/- mice, overactivation of PDGFRβ signaling slightly diminished fibrous cap area. These data suggest that in plaques with elevated PDGFRβ activation, VSMC apoptosis counterbalances or even exceeds VSMC proliferation and migration. Risk factors like hyperglycemia can further accelerate VSMC cell death. A recent study showed that hyperglycemia advances vascular cell death by blunting PDGF-dependent anti-apoptotic signaling.^39^ Our further *in vitro* analysis revealed that PDGFRβ-induced activation of PI3K led to c-Myc dependent downregulation of Bcl-2, increasing VSMĆs susceptibility to apoptosis. C-Myc expression is suppressed by transcription factor FOXO1,^40^ which is inactivated by PI3K/AKT dependent phosphorylation.^41^ Hence, activation of PI3K/AKT induces c-Myc via inactivation of FOXO1. MYC has been shown to repress both mRNA and protein expression of pro-survival members such as BCL- 2 or BCL-XL.^35^ Consistently, Bcl-2 expression is markedly reduced in plaques from LDLR^-/-^smLRP1^-/-^ mice. In human atherosclerotic lesions, VSMC apoptosis is notably increased, and apoptotic cells mostly lack Bcl-2.^42^ PDGFRβ expression and activation have been found at increased levels.^43^ Furthermore, VSMCs derived from human atherosclerotic plaques are more sensitive to apoptosis^44^ and display enhanced expression of c-Myc^36,45^ compared to cells derived from non-diseased vessels. Expression profiling and quantitative proteomics analysis of PDGF-treated human VSMCs identified c-Myc as the transcription factor most significantly networked with up-regulated genes.^46^ These interesting correlations suggest that elevated levels of PDGF and PDGFRβ in diseased vessels might increase the susceptibility of VSMCs to apoptosis via c-Myc dependent downregulation of Bcl-2 and thereby promotes growth and destabilization of atherosclerotic plaques.

These data appear unexpected as many previous publications suggest an exclusive anti-apoptotic effect of PDGF *in vitro* and *in vivo.*^1,15,21^ However, the present data demonstrate that this effect is time-dependent and that in addition to these survival effects, a pro-apoptotic pathway arises in parallel. Our data provides evidence that like anti-apoptotic PDGF signaling, this pro-apoptotic pathway also exclusively depends on PI3K. Analysis of chimeric c-fms/PDGFRβ (ChiR) mutants revealed that deletion of the PI3K binding site in the PDGFRβ polypeptide abrogated this effect, whereas activation of PI3K alone was sufficient to induce apoptosis to a similar extent than ChiR-WT. Interestingly, activation of the F771 mutant which lacks the binding site for the negative regulator of Ras, RasGAP, mediated a much more pronounced increase of apoptosis than the ChiR-WT. This was expected since accumulation of GTP-bound Ras is necessary to fully activate PI3K.^47^ A Ras/ERK dependent effect in this context appears unlikely since inhibition of MEK1/2 upstream of ERK was ineffective. Whether PDGF/PI3K-induced survival or pro-apoptotic effects prevail in VSMCs, seems to depend on several parameters including the time interval following PDGFRβ activation, additional extracellular stimuli, the cell type, and differential activation of diverse PI3K isoforms. Interestingly, selective inhibition of catalytic PI3K subunits indicated isoform specific effects on PDGF-mediated apoptosis. While selective inhibition of PI3K/p110α did not affect long-term PDGF dependent apoptotic effects on H_2_O_2_ treated cells, short-term survival effects of PDGF in VSMCs primarily depended on p110α. Conversely, inhibition of p110δ and p110β rather ameliorated cell death without affecting PDGF-dependent survival effects. These findings suggest that PI3K isoform p110α mediates survival effects on VSMCs, whereas p110δ and p110β are rather responsible for apoptotic signaling under chronic PDGF conditions. This data is consistent with our previous finding demonstrating that under short term conditions, only p110α is recruited to the activated PDGFRβ.^48^ Accordingly, stimulation with PDGF and downstream activation of PI3K elicit only transient survival effects, followed by a delayed but persistent downregulation of Bcl-2 favouring apoptosis.

When Akt signaling is shut off, and Bcl-2 expression is low, celĺs fate depends on the extracellular context. In advanced atherosclerotic plaques harbouring a pro-inflammatory and pro-apoptotic milieu, PDGF-dependent downregulation of Bcl-2 accelerates VSMC apoptosis induced by components like tumour necrosis factor α, Fas-L, and oxidised low-density lipoprotein. Other stimuli can directly modulate PDGF-dependent apoptosis or survival signaling. Interleukin-1β prolongs PDGF-dependent Akt activation, thereby promoting cellular survival^34^. Finally, PDGF-induced survival versus pro-apoptotic effects appear cell type-dependent. In fibroblasts, chronic stimulation with PDGF increases apoptotic cell death suggesting that this effect is not restricted to VSMCs.^49^ However, chronic PDGF stimulation does not exert pro-apoptotic effects on cardiomyocytes.^50^ Since cardiomyocytes in contrast to VSMCs or fibroblasts are not able to proliferate upon mitogenic stimulation, one might speculate that PDGF-mediated proliferation and apoptosis are tightly coupled. This is supported by the fact that PDGFRβ-induced pro-apoptotic and proliferative effects are mediated by the same proteins: PI3K and c-Myc.^21,46^ Therefore, this newly identified PDGF driven pro-apoptotic signaling pathway in VSMCs may provide a failsafe program to limit uncontrolled cell growth under inappropriate conditions. In this context, it is important to note that VSMCs are highly plastic. Embedded within the aorta tunica media in their contractile state, VSMCs modulate pressure and blood flow. When stimulated by diverse growth factors and cytokines, contractile VSMCs de-differentiate, and enter a highly plastic state. PDGF suppresses contractile markers, including MYH11 and ACTA2. Upon de-differentiation, several stimuli drive VSMC phenotypes toward multiple distinct cellular fates.^19^ Therefore, one might speculate that PDGF exerts pro- or anti-apoptotic effects depending on the VSMC phenotype, thereby promoting or suppressing distinct VSMC clones. However, under advanced pathophysiological conditions in atherogenesis this failsafe program might outnumber cellular proliferation leading to unmeant consequences like thinning of the fibrous cap, plaque rupture, and finally myocardial infarction or stroke.

In conclusion, our data demonstrate that in addition to PDGF acting as a cellular survival factor, chronic PDGF/PI3K signaling may lead to pro-apoptotic effects on VSMCs via regulation of Bcl-2 and c-Myc, and pro- versus anti-apoptotic PDGFRβ signaling is differentially regulated via catalytic class IA PI3K isoforms. The relative contribution of PDGF-dependent proliferation/migration versus VSMC apoptosis during long-term PDGFRβ activation crucially determines plaque composition during atherogenesis, resulting in apoptosis and lower number of VSMCs in the fibrous cap and typical features of plaque instability.

## Acknowledgements

The technical support by Frank Oberhäuser and Maximilian Becker is greatly acknowledged. This works contains part of a doctoral thesis by MS.

