## Supplementary Material for "An Apoptotic Failsafe Program Involving PDGF-dependent PI 3- Kinase Activity Contributes to Plaque Composition in Atherosclerosis"

### Supplemental Material

**Supplemental Figure 1:** Pre-stimulation with PDGF dose dependently increases $H_2O_2$-induced apoptosis.

**Supplemental Figure 2:** Schematic diagram illustrating the cytoplasmic domains of chimeric c-fms/PDGFR (ChiR) mutants used in this study.

**Supplemental Figure 3:** Long-term stimulation with PDGF decreases Bcl-2 expression.

**REFERENCES**

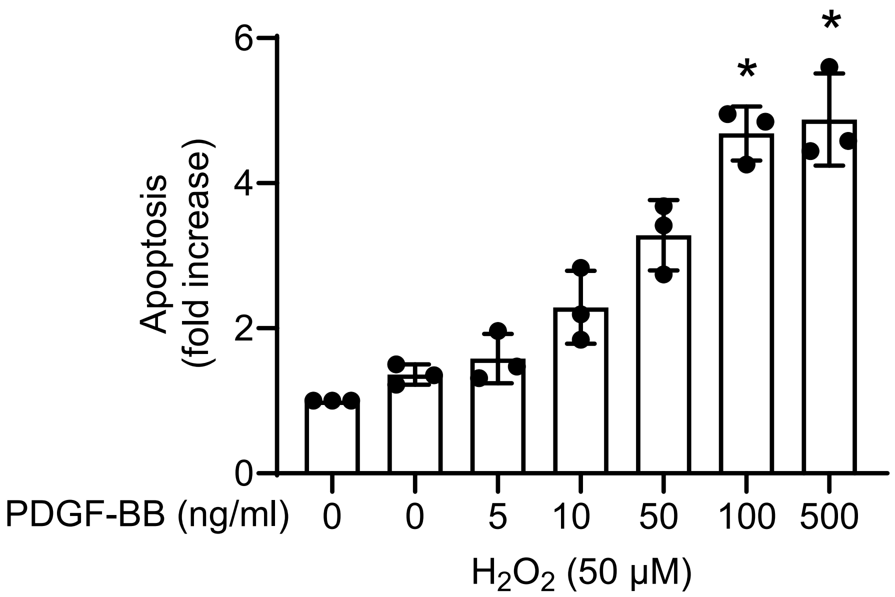


**Supplemental Figure 1** Prestimulation with PDGF dose dependently increases $H_2O_2$-induced apoptosis. Analysis of $H_2O_2$-induced DNA fragmentation in cells pretreated with the indicated concentrations of PDGF. n = 3. Data is presented as means ± SD. *P* values were determined by Repeated-Measures ANOVA followed by Tuckey´s multiple comparison. (*$p$<0.05).

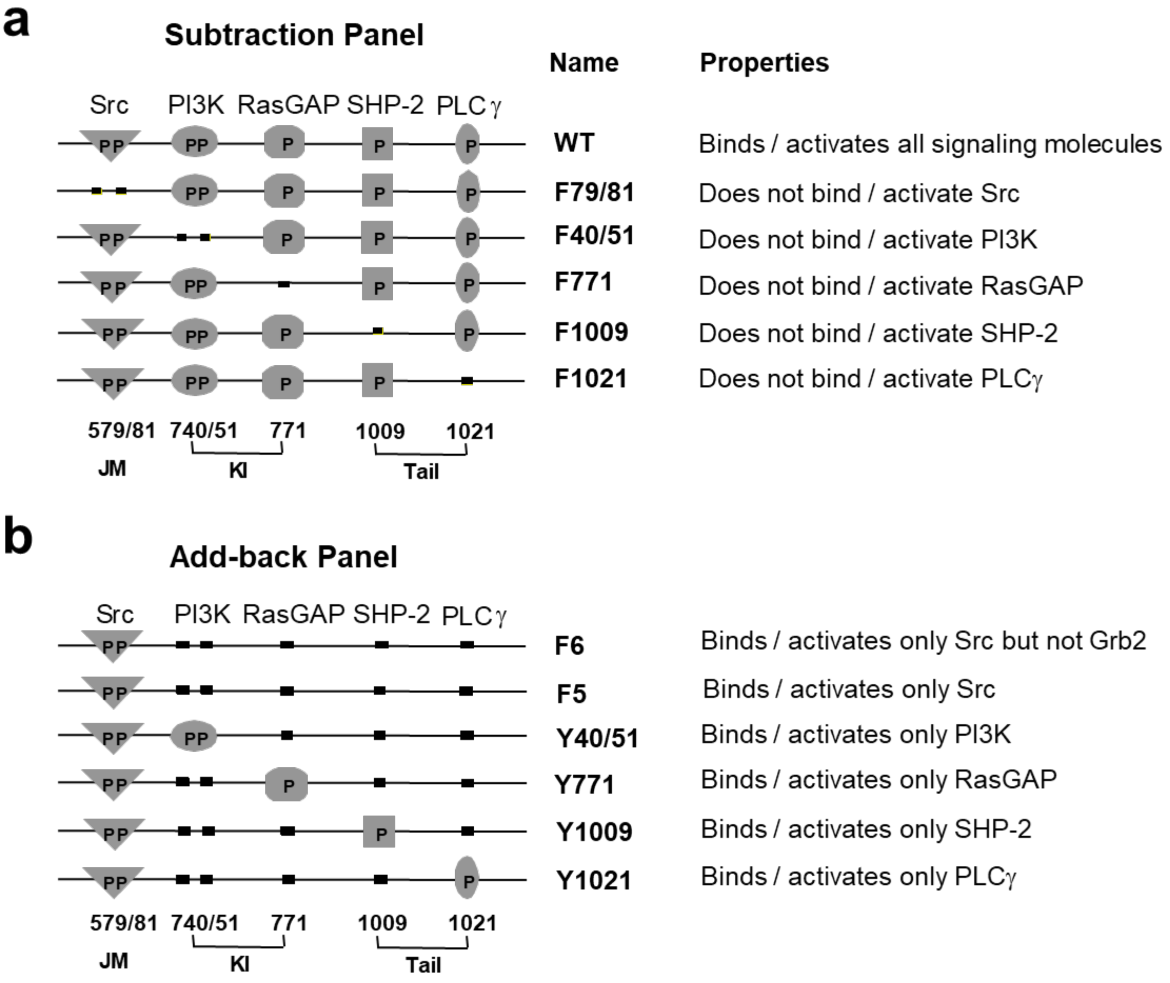


**Supplemental Figure 2** Schematic diagram illustrating the cytoplasmic domains of chimeric c-fms/PDGFR (ChiR) mutants used in this study[1]. Tyrosine phosphorylation sites are represented as *P*, and Tyr to Phe substitutions are indicated as black squares. Signaling enzymes which stably associate with the receptor mutants are are indicated at the *top* of the schemes. The nomenclature of the "subtraction panel" (a) and "add-back panel" (b) of the ChiR is quoted to the *right*. In the subtraction panel, the names indicate which of the tyrosine residues have been replaced with phenylalanine, and in the add-back panel the name of each mutant denotes which of the mutations in the F5 construct has been restored. Tyr 579 and 581 are located in the juxtamembrane domain (*JM*) of the receptor and are required for Src binding; Tyr 741, 752, and 771 are in the kinase insert (*KI*) of the receptor and are responsible for PI3K and RasGAP binding, respectively; Tyr 1009 and 1021 are located in the C-terminal *tail* region of the receptor, and are involved in the binding of SHP-2 and PLCγ, respectively. The F6 construct contains an additional mutation at the Grb2 binding site Tyr 716.

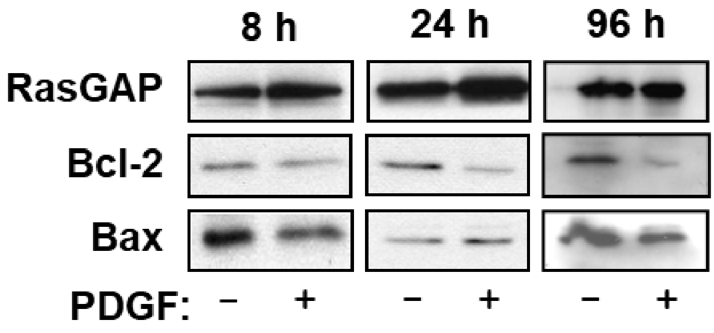


**Supplemental Figure 3** Long-term stimulation with PDGF decreases Bcl-2 expression. Representative immunoblots with lysates from VSMC treated for the indicated periods of time with PDGF and probed with antibodies against Bcl-2, Bax, and RasGAP (loading control).
